# Providing new insights on the byphasic lifestyle of the predatory bacterium *Bdellovibrio bacteriovorus* through genome-scale metabolic modeling

**DOI:** 10.1101/2020.01.13.904078

**Authors:** C Herencias, MA Prieto, J Nogales

**Affiliations:** Microbial and Plant Biotechnology Department, Biological Research Center, CSIC, Madrid, Spain.; Interdisciplinary Platform for Sustainable Plastics towards a Circular Economy-Spanish National Research Council (SusPlast-CSIC), Madrid, Spain.; Department of Systems Biology, Centro Nacional de Biotecnología, CSIC, Madrid, Spain

## Abstract

In this study we analyze the growth-phase dependent metabolic states of *Bdellovibrio bacteriovorus* by constructing a fully compartmented, mass and charge-balanced genome-scale metabolic model of this predatory bacterium (*i*CH457). Considering the differences between life cycle phases driving the growth of this predator, growth-phase condition-specific models have been generated allowing the systematic study of its metabolic capabilities. Using these unprecedented computational tools, we have been able to analyze, from a system level, the dynamic metabolism of the predatory bacteria as the life cycle progresses. We provide solid computational evidences supporting potential axenic growth of *B. bacteriovorus*’s in a rich medium based its encoded metabolic capabilities. Our systems-level analysis confirms the presence of “energy-saving” mechanisms in this predator as well as an abrupt metabolic shift between the attack and intraperiplasmic growth phases. Our results strongly suggest that predatory bacteria’s metabolic networks have low robustness, likely hampering their ability to tackle drastic environmental fluctuations, thus being confined to stable and predictable habitats. Overall, we present here a valuable computational testbed based on predatory bacteria activity for rational design of novel and controlled biocatalysts in biotechnological/clinical applications.

**AUTHOR SUMMARY:** Bacterial predation is an interspecific relationship widely extended in nature. Among other predators, *Bdellovibrio* and like organism (BALOs) have received recently a great attention due to the high potential applications such as biocontrol agent in medicine, agriculture, aquaculture and water treatment. Despite the increasing interest in of this predatory bacterium its complex lifestyle and growth conditions hamper the full exploitation of their biotechnological properties. In order to overcome these important shortcomings, we provide here the first genome-scale model of a predator bacterium constructed so far. By using the model as a computational testbed, we provide solid evidences of the metabolic autonomy of this interesting bacterium in term of growth, as well as its dynamic metabolism powering the biphasic life style of this predator. We found a low metabolic robustness thus suggesting that Bdellovibrio is more niche-specific than previously thought and that the environmental conditions governing predation may be relatively uniform. Overall, we provide here a valuable computational tool largely facilitating rational design of novel and controlled predator-based biocatalysts in biotechnological/clinical applications.

## INTRODUCTION

Predation is a biological interaction where an individual, the predator, feeds on another, the prey, to survive. Since predation has played a central role in the diversification and organization of life, this system provides an interesting biological model from both an ecological and evolutionary point of view. Predation is an example of coevolution where the predator and prey promote reciprocal evolutionary responses to counteract the adaptation of each other [1]. This interspecific relationship is widely extended in nature, including the microbial world where the main predators are bacteriophages, protozoa and predatory bacteria [2]. Focusing on bacteria, this group is composed, among others, by *Bdellovibrio* and like organisms (BALOs) which are small, highly motile, and aerobic gram-negative predatory bacteria that prey on a wide variety of other gram-negative bacteria. Originally discovered in soil [3], BALOs are ubiquitous in nature. They can be found in terrestrial and aquatic habitats, bacterial biofilms, plants, roots, animals and human feces [4] and lung microbiota [5]. *B. bacteriovorus* is the best characterized member of the group of BALOs and the genome of different strains, including HD100, Tiberius and 109J have been sequenced providing a reliable source of genetic information [6–8].

*B. bacteriovorus* exhibits a biphasic growth cycle (Fig 1), including a free-swimming attack phase (AP) and an intraperiplasmic growth phase (GP) inside the preýs periplasm forming the so-called bdelloplast structure. During AP, free living cells from extracellular environment are in active search for new preys. After attachment, and once the predator-prey interaction is stable and irreversible, the predator enters in the prey’s periplasm, where it grows and replicates DNA during the GP using the cytoplasm of the prey cell as a source of nutrients and biomass building blocks. When the prey is exhausted, *B. bacteriovorus,* grown as a filament, septs into several daughter cells and, through its large array of hydrolytic enzymes, lyses the ghost-prey’s outer cell membrane and releases into the medium [9]. Interestingly, host-independent (HI) mutants of *Bdellovibrio* strains have been developed under laboratory conditions. These HI predators are able to grow axenically (without prey) in a rich-nutrient medium mimicking the dimorphic pattern of elongated growth, division and differentiation [10]. It is worth noticing that the axenic growth of these mutant strains is given by a mutation in the host interaction (hit) locus, which has been described as being involved in regulatory and/or scaffold elements [11]. This argues in favor of this mutation having no metabolic (enzymatic) impact. In fact, the main metabolism of these HI derivatives should not have suffered changes with respect to the wild type *Bdellovibrio* strains.

**Fig 1.**
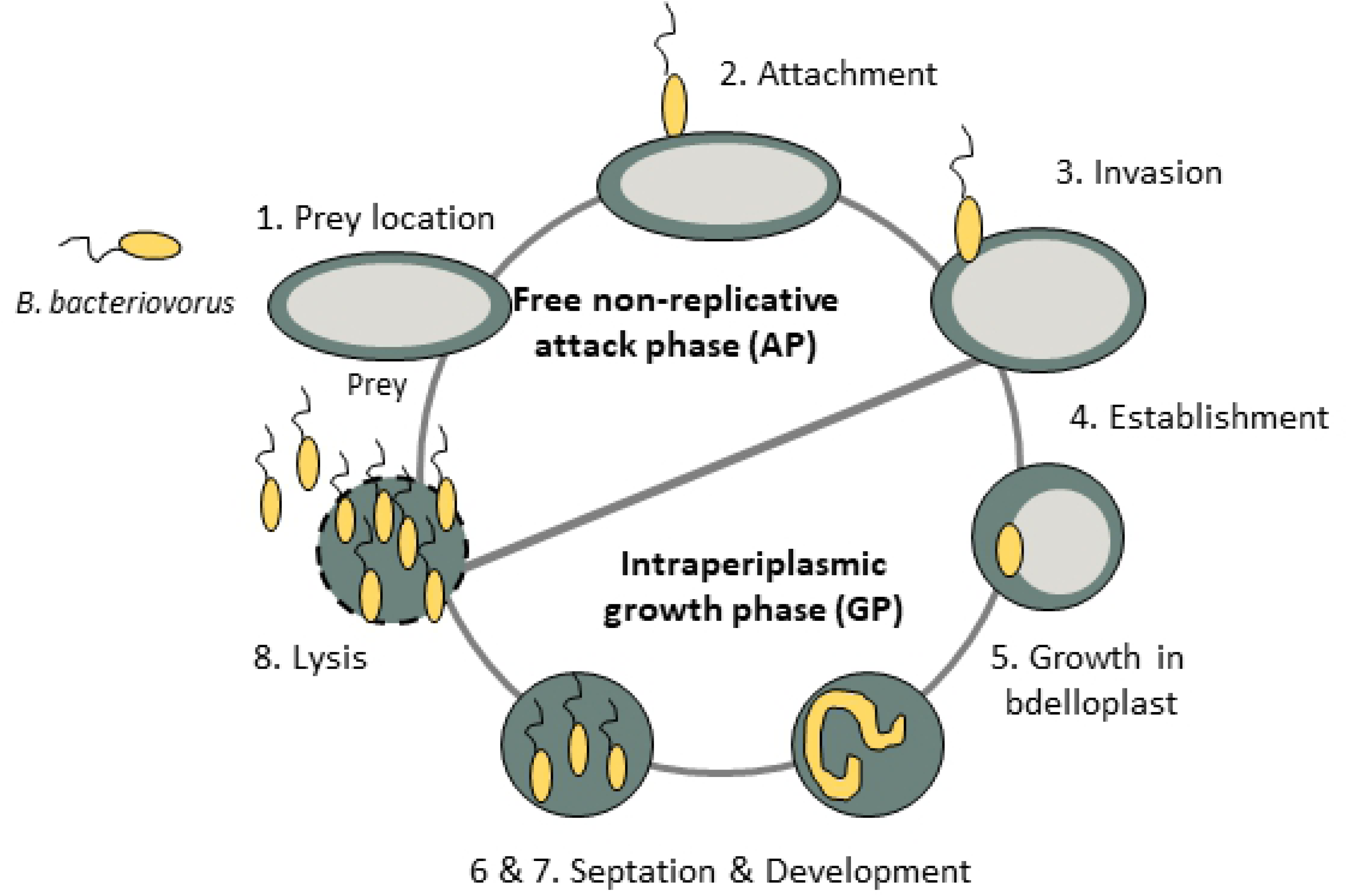
Lifecycle of *Bdellovibrio bacteriovorus* HD100. 1) Prey location: *B. bacteriovorus* moves towards prey-rich regions. 2) Attachment: the predator anchors to the host cell, which leads to the infection. 3) Invasion: *B. bacteriovorus* enters the periplasm of the prey cell. 4 and 5) Growth in bdelloplast and development: the prey has a rounded appearance due to cell wall modification and *B. bacteriovorus* grows in the periplasm and replicates its DNA. *B. bacteriovorus* uses the prey cytoplasm as a source of nutrients. 6 and 7) Septation and development: the predator septs when resources become limited and it matures into individual attack phase cells. 8) Lysis: mature attack-phase cells lyse the cell wall of the bdelloplast, initiating the search for fresh prey. The complete cycle takes about 4 h.

*B. bacteriovorus*’ extraordinary repertoire of susceptible preys allows for a wide range of potential applications based on its predatory capability, such as biocontrol agent in medicine, agriculture, aquaculture and water treatment [12–15]. Furthermore, it has been proposed as an excellent source of valuable biotechnological enzymes and as a biological lytic tool for intracellular products, due to its hydrolytic arsenal [4,16,17]. Moreover, regarding its unique lifestyle it represents a good model for evolution studies focusing, for example, on the origin of the eukaryotic cell [18, 19]. Despite the interest that this predatory bacterium’s potential applications have recently aroused among the scientific community, its complex lifestyle and growth conditions make it hard to implement metabolic and physiological studies. As a direct consequence, to date, its physiology and metabolic capabilities remain an enigma to a large extent [20]. Moreover, the potential of this predator to be used as a biotechnological chassis depends on the quantity and quality of the available metabolic knowledge. Therefore, expanding the knowledge of this predatory bacterium as a shuttle for the further full exploitation of its unique biotechnological applications would require a reliable platform driving the rational understanding of its characteristics.

Following this aim, the advent of genomic age and the subsequent large amount of derived high-throughput data, have largely contributed to deeper understanding of microbial behavior, at system level [21]. Specifically, genome-scale metabolic models (GEMs) are being used to analyze bacterial metabolism under different environmental conditions [22, 23]. GEMs are structured representations of the metabolic capabilities of a target organism based on existing biochemical, genetic and phenotypic knowledge which can be used to predict phenotype from genotype [24].

The application of Constraint-Based Reconstruction and Analysis (COBRA) approaches [25] together with specific GEMs have been successfully applied for better understanding of interspecies interactions such as mutualism, competition and parasitism providing important insights into genotype-phenotype relationship [26]. Despite GEMs being powerful tools to elucidate the metabolic capabilities of single systems, addressing the complex metabolism of bacterial predators having biphasic growth-cycles such as *B. bacteriovorus* is challenging and has remained elusive so far.

We provide here the first step toward the metabolic understanding at system level of *B. bacteriovorus* by the reconstruction of its metabolism at genome-scale. We further use this cutting edge computational platform as a test bed for the integration and contextualization of transcriptomic and physiological data shedding light on the biphasic lifestyle of this predatory bacterium.

## RESULTS

### Characteristics of *B. bacteriovorus* metabolic reconstruction

A genome-scale metabolic model (*i*CH457) including the metabolic content derived from genome annotation and available biochemical information was created for *B. bacteriovorus* HD100. *i*CH457 does not differentiate AP and GP, but it is a powerful tool for determining and analyzing the potential metabolic capabilities of the system from a global perspective. All the gene-protein-reaction associations (GPRs) included in the model were subject to a rigorous manual curation process in order to ensure the quality of the final model (Fig. 2). Several open reading frames (ORFs) were annotated de novo and/or re-annotated during the reconstruction process. For instance, from the initial 75 ORFs included in the reconstruction draft belonging to amino acid metabolism based on bioinformatics evidence, only 65 (87%) were finally included according to bioinformatics and literature-based evidences. Moreover, during the manual curation process, we confirmed (by sequence homology) the function of several genes related to amino acids metabolism and the hydrolytic enzymes involved (Tables S4 and S5). For instance, gene *bd0950*, annotated as an unspecific acetyltransferase, was specifically associated with an UDP 2,3-diamino-2,3-dideoxy-D-glucose acyltransferase, while gene *bd2095*, wrongly annotated as encoding for an acetyl-CoA C-acetyltransferase, was unequivocally re-annotated as a 3-ketoacyl-CoA thiolase.

**Fig 2.**
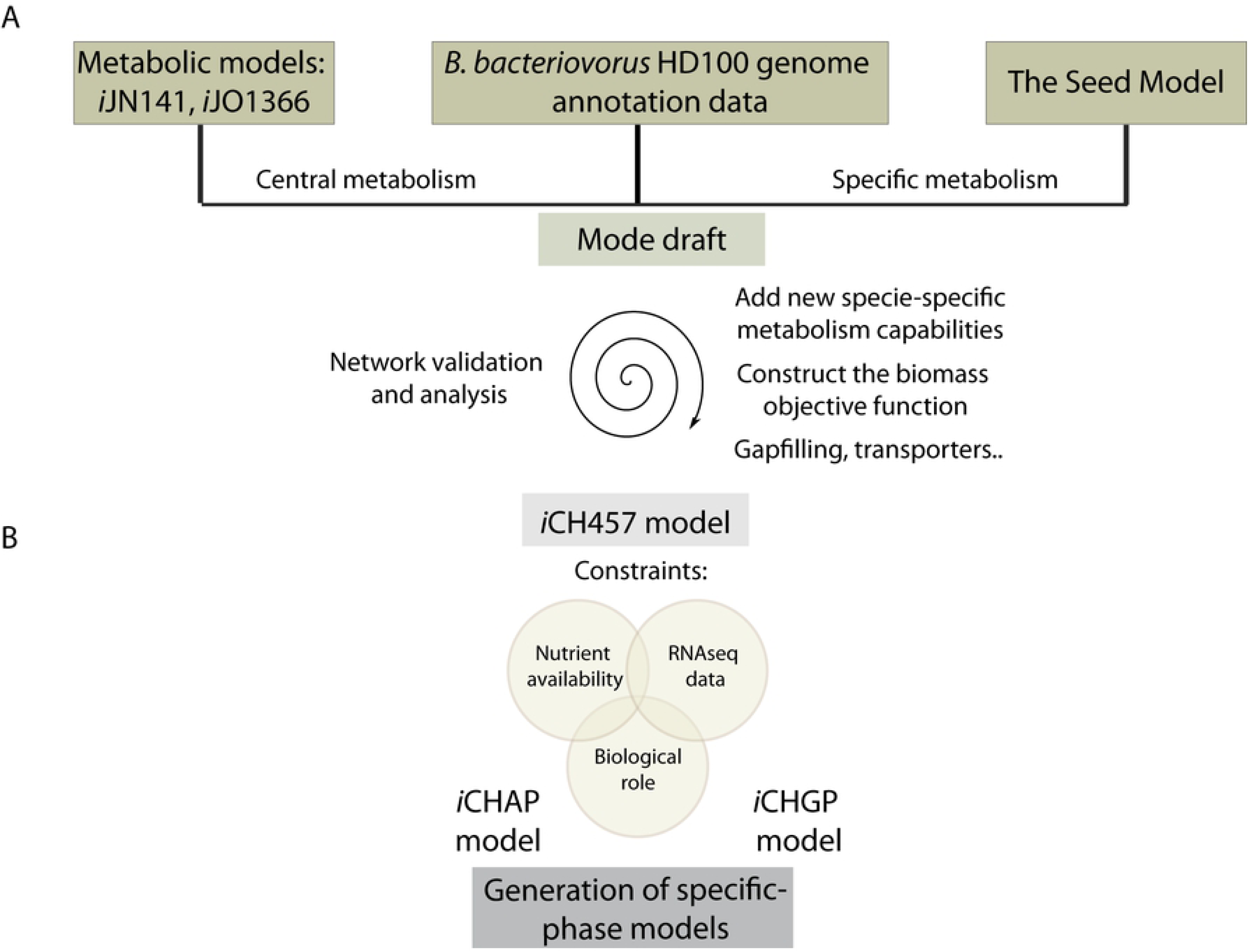
*i*CH457 metabolic model pipeline. **A)** The draft of metabolic reconstruction was based on available metabolic models (*i*JN1411 and *i*JO1366), the genome sequence of *B. bacteriovorus* HD100 and the automatic model Seed. Manual curation is required to accurately fine-tune the information contained in the metabolic model and several steps of network validation and analysis are required to finally obtain the metabolic model *i*CH457. **B)** Generation of condition-specific models: *i*CHAP and *i*CHGP. The general model *i*CH457 was constrained based on nutrient availability (minimal and rich *in silico* media), biological role (ATP production or biomass generation) and transcriptomic available data (Karunker et al., 2013)*. GIM^3^E algorithm was used to construct the condition-specific models.

*i*CH457 includes 457 ORFs, which represent 13 % of the coding genes in the genome, whose gene products account for 705 metabolic and transport reactions (accounting for 70.5 % of the model’s total reactions). The model was completed with the inclusion of 296 non-gene associated reactions (29.5 %) based on physiological and/or biochemical evidences supporting their presence in *B. bacteriovorus*. For instance, reactions related to the ACP acyltransferase (G3PAT) needed for glycerophospholipid biosynthesis were included based on the physiological evidence provided by Nguyen and col. and Muller and col. [27, 28]. Overall, *i*CH457 accounts for a total of 1001 reactions and 955 metabolites distributed in three different compartments: cytoplasm, periplasm and extracellular space.

Reactions from *i*CH457 fall into 12 main functional categories (Fig 3). It is noteworthy that cell envelope metabolism seems to be the most represented group with a total of 222 reactions. In this important group we found reactions involved in the metabolism of peptidoglycans, lipopolysaccharides, glycerophospholipids, and murein. Across this group, catabolic reactions including reactions involved in the degradation of peptidoglycan by specific carboxypeptidases represent up to 37%. This high number of hydrolytic reactions present in *i*CH457 is consistent with the important role of these enzymes in the degradation of the prey’s cell wall to penetrate into the periplasm, completion of the growth cycle and also in recycling the envelope components [29].

**Fig 3.**
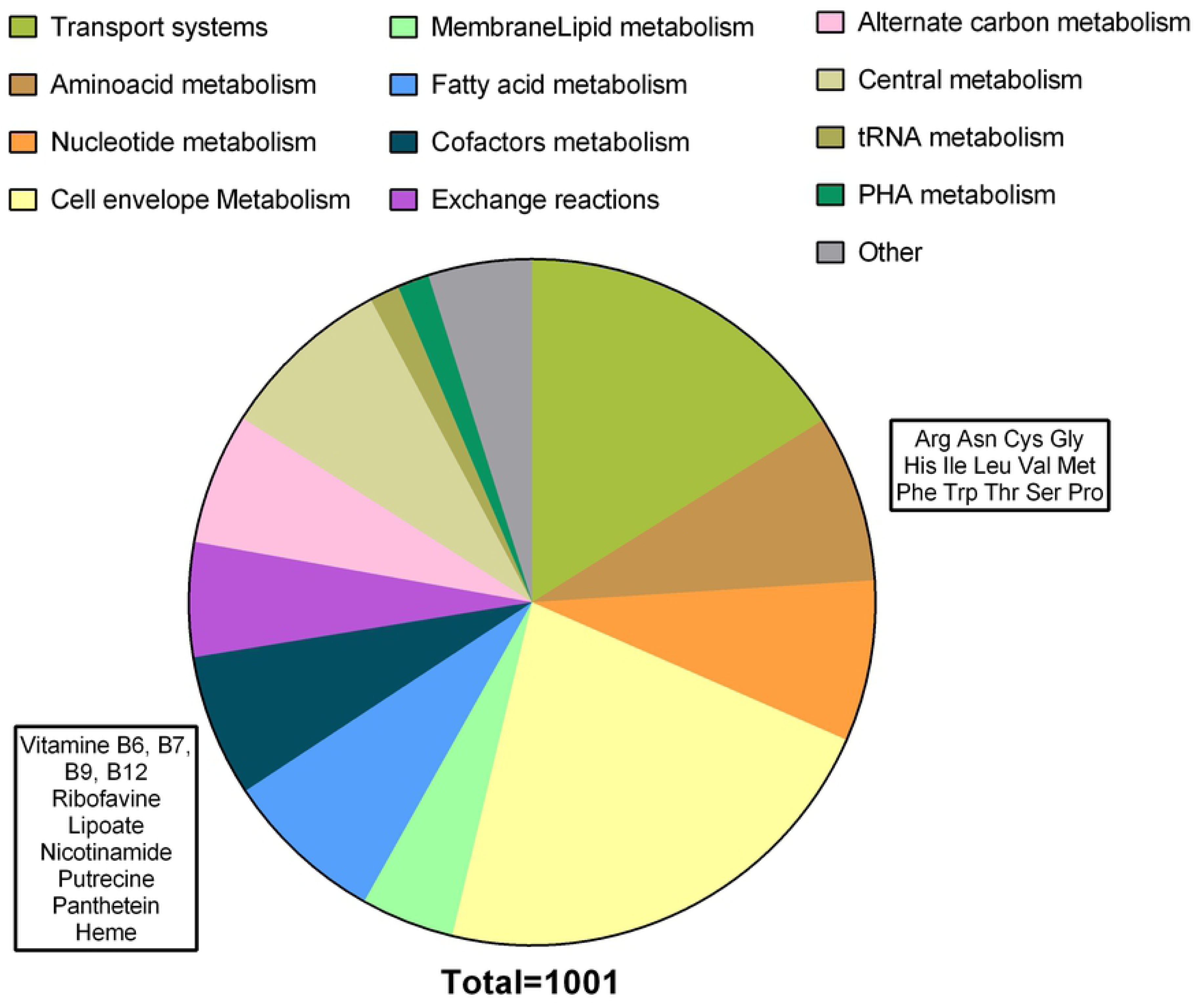
Distribution of the reactions of the system in 12 global functional categories. The metabolites inside the rectangles correspond with the auxotrophies in the cofactor metabolism (dark blue fraction) and the amino acid metabolism (brown fraction), respectively. Asterisks (*) show the metabolite groups that contain auxotrophies.

In the past 15 years, GEMs have garnered considerable research attention and numerous metabolic reconstructions have been generated for several organisms [30]. The metabolic models within the δ-proteobacteria group are underrepresented among this phylum and only a few of them have been constructed, for instance for *Geobacter* spp. and *Desulfovibrio vulgaris* [31–33]. Thus, the model of *B. bacteriovorus* HD100, *i*CH457, represents a new model within this group, which, as depicted in Table 1, provides a complete reconstruction of this important bacterial group in terms of the metabolites and reactions included. Furthermore, other microbial interactions have been recently modeled, such as the syntrophic association between *Desulfovibrio vulgaris* Hildenborough and *Methanococcus maripaludis* S2 [32]. The metabolic reconstruction of this interaction highlighted the potential use of *in silico* predictions to capture growth parameters and community composition of bacterial communities. However, this is the first metabolic model of a predatory bacterium.

**Table 1.** Comparison of the metabolic properties of *i*CH457 compared with other δ-proteobacteria (*Geobacter* spp. and *Desulfovibrio vulgaris*) metabolic models and with the well-establish metabolic reconstruction of *P. putida* (*i*JN1411) and *E. coli* (*i*JO1366).

|  | *iJN1411* | *iJO1366* | *Geobacter metallireducens* | *Geobacter sulfirreducens* | *Desulfovibrio vulgaris* Hildenborough | *iCH457* |
| --- | --- | --- | --- | --- | --- | --- |
| Protein-coding Genes | 5350 | 4405 | 3532 | 3530 | 3379 | 3584 |
| Genes (%) | 1411<br>(26%) | 1366<br>(31%) | 747 (21%) | 588 (17%) | 744 (22%) | 457 (13%) |
| Metabolites | 2057 | 1136 | 769 | 541 | 1016 | 956 |
| Reactions | 2754 | 2251 | 697 | 523 | 951 | 1001 |
| Reference | (Nogales et al., 2017) | (Orth et al., 2011) | (Sun et al., 2009) | (Mahadevan et al., 2006) | (Flowers et al., 2018) | This work |

### Model-driven assessment of auxotrophies and biomass building block transport systems highlight the predatory lifestyle of *B. bacteriovorus*

During the reconstruction process, we identified several incomplete biosynthetic pathways (amino acids, cofactors, and vitamins) that are consistent with the numerous auxotrophies previously reported for strain HD100 [6]. However, model-based analyses provide an integrated overview of the complete metabolic network of this predator, including metabolic gaps potentially responsible for the auxotrophies. This is because instead of analyzing just the main synthetic pathways, such *in silico* analyses consider the global metabolism, including alternative and/or secondary/accessory biosynthetic routes.

In fact, model-based analyses identified up to 24 different auxotrophies. For instance, from the 20 proteinogenic amino acids, external supply of 14 of them was required to achieve *in silico* growth, including arginine, asparagine, cysteine, glycine, histidine, methionine, leucine, isoleucine, valine, phenylalanine, tryptophan, threonine, serine and proline. In addition, external supply of several cofactors including riboflavin, nicotinamide, putrescine, folic acid, pantothenate, pyridoxal phosphate, biotin and lipoate was needed in order to achieve *in silico* growth (Fig 2A).

Concerning nucleosides monophosphate, we found that *B. bacteriovorus* has the ability to synthetize *de novo* all these key biomass building blocks despite nucleosides derived from the hydrolysis of prey having been traditionally suggested as a source of nucleic acids [34]. Supporting this computational analysis, radiotracer studies showed that strain 109J mainly utilized host nucleosides monophosphate during intraperiplasmic growth, however it was also able to synthesize its own pool of nucleotides [35, 36]. This phenomenon has been traditionally explained in the context of an “energy-saving” mechanism. Similarly, this mechanism has also been reported, and validated by our *in silico* analysis, for phospholipid assimilation and the recycling of some unaltered or altered fatty acids from the prey. Thus, while model analysis confirmed a complete and likely functional fatty acid *de novo* biosynthetic pathway, the direct uptake of these biomass building blocks has been largely reported. This demonstrates a more efficient incorporation of cellular components from the prey [27, 37].

Due to their lifestyle and the obligate requirement of obtaining essential biomass building blocks from prey, the transport subsystem became an important key for the survival of *B. bacteriovorus*. In fact, this category was found to be one of the most representative in terms of number of reactions (161), highlighting their importance in cellular interchange. Although a comprehensive analysis of the transport systems in the predator has been previously reported [38], the predicted substrate specificity needs more experimental support. Remarkably, *i*CH457 model accounts for 67 % of the annotated transport system reported in the genome.

It is worth emphasizing the case of peptide transporters; despite amino acids from protein breakage having been suggested as major carbon and energy sources during the intraperiplasmic growth of *B. bacteriovorus* [39], we noticed a significant lack of specific amino acid transporters during the model reconstruction and functional validation process. Instead, we found a large number of di- and tripeptide transporters, suggesting that the predator might be taking up small peptides from the prey.

Overall, model-based analyses largely supported the presence of energy-saving mechanisms in *B. bacteriovorus* targeting the biosynthesis of nucleotide monophosphates and phospholipids, but not of amino acids or vitamins whose availability depends exclusively on the prey. Likewise, detailed analysis of transport systems included in the model suggests *B. bacteriovorus*’ ability to obtain oligopeptides through prey proteins cleavage and use them as its main source of carbon, nitrogen and energy during GP.

### *i*CH457 exhibits high accuracy predicting physiological states of *B. bacteriovorus* under different nutrients scenarios

A model’s capability of providing accurate predictions of empirically-supported knowledge of a target organism’s functional states is a key feature in order to assess the accuracy and completeness of the final reconstruction. However, the obligate predatory lifestyle of *B. bacteriovoru* and the complex environment provided by the prey, in terms of nutrients, prove challenging when using classical validation workflows based on single nutrient sources. Therefore, for *i*CH457 validation, we took advantage of spontaneous HI *Bdellovibrio* strains developed under laboratory conditions. Such HI strains exhibit a similar lifecycle when growing in a rich medium to the wild type strain growing inside the intraperiplasmic space of the prey [40]. Indeed, because the HI phenotype has been attributed to putative regulatory and/or scaffold mechanisms rather than to metabolic genes (enzymes) [41, 42], these HI strains are supposed to possess metabolic capabilities identical to those of the parental strains. Thus, for the GEMs validation process, including potential carbon sources and biomass generation rates, we decided to use data from HI *Bdellovibrio* strains for *i*CH457 validation.

Specifically, we validated the predictive capabilities of the *i*CH457 by comparing *in silico* results with experimentally determined biomass production and growth rates of the HI strain *B. bacteriovorus* 109 Davis [43]. The *in silico* growth rates were calculated using minimal medium supplemented with selected carbon sources (S1 Text). *i*CH457 was very precise predicting growth rate on five different carbon sources, with an accuracy close to 100% in the case of glutamate, glutamine and succinate, and 70% for pyruvate and lactate (Fig 4A). The slight discrepancies found between *in silico* predictions and *in vivo* results might be explained by an incomplete formulation of biomass function or higher energy maintenance requirements under the simulated conditions not accounted for in the current reconstruction. In addition, higher *in silico* growth rates are often found due to the intrinsic nature and limitations of FBA. FBA presumes a final evolution state in stark contrast with the potential scenario found *in vivo* which could lack the proper adaptation to these metabolites as a primary carbon source [44]. Also, FBA only predicts steady-state fluxes and does not account for any regulatory constraints, which should play an important role in the uptake of substrates from the extracellular medium [45]. Overall, our model predictions showed a significant accuracy that is comparable to other high-quality genome-scale models already available [46].

**Fig 4.**
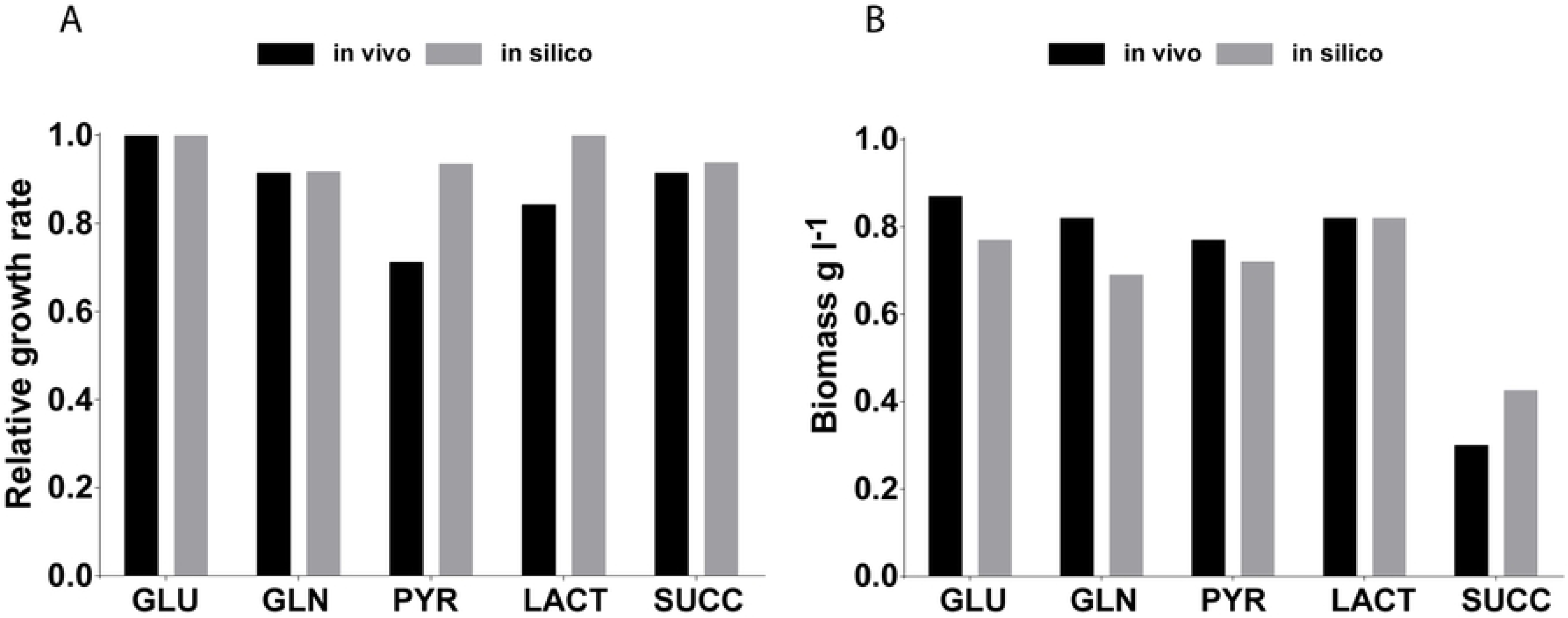
Evaluation of the metabolic capabilities of *i*CH457. **A) Comparison of the growth performance of the *in silico i*CH457 strain and a derivative strain of *B. bacteriovorus* 109 Davis on different carbon sources.** Experimental values of growth rate were calculated using the mass doubling time previously compiled in Ishiguro et al. (Ishiguro, 1974). The *in silico* growth rate was calculated with the minimal medium defined in Annex 4 supplemented with the tested carbon source. B) Comparison of the biomass production predicted *in silico* with the available experimental data performed with the prey-independent *B. bacteriovorus* 109 Davis (Ishiguro, 1974). *In vivo* and *in silico* biomass data were expressed as Kendall’s rank correlation coefficient (τ = 0,88) for *i*CH457. GLU: glutamate, GLN: glutamine, PYR: pyruvate, LACT: lactate, SUCC: succinate.

Beyond the availability to predict growth rates, it is valuable to assess the model’s ability to predict the maximum amount of biomass produced from known concentrations of given carbon and energy sources. Similar high accuracy was found regarding the predictability of biomass production between *in silico* and experimental data (Kendall’s coefficient τ = 0.88) (Fig 4B). It is noteworthy that the *in silico* analysis provided in these evaluations largely confirmed the prey-independent metabolic states, thus shedding light on the predator’s potentially autonomous metabolism. These results are in good agreement with the large amount of HI derivative strains isolated previously [47] and the recent description of the metabolic response of AP cells in NB medium to synthesize and secrete proteases [48]. Therefore, the obligate predatory lifestyle of *B. bacteriovorus* should be questioned, at least from a metabolic point of view.

Overall, the high accuracy exhibited by *i*CH457 encourages us to use the model to characterize and better understand the metabolic states that underline the biphasic growth cycle of *B. bacteriovorus*.

### Reaction essentiality towards understanding the predator’s lifestyle

It is well-known that the environmental conditions and natural habitat of a given bacterium largely influence its evolutionary traits, including processes of genome expansion/reduction. Therefore, and taking advantage of *i*CH457, it would be interesting to address from a computational perspective whether the genome content of the predator has been influenced by its complex lifestyle. We identified a set of essential reactions in *i*CH457. The network reaction(s) associated with each gene was individually “deleted” by setting the flux to 0 and optimizing for the biomass function. A reaction was defined as essential if after constrained it the growth rate decreased to less than 10% of wild type model. To properly contextualize the reaction essentiality analysis, we compared our results with those from some free-living organisms such as *P. putida* KT2440 (*i*JN1411) [46], *E. coli* strain K-12 MG1655 (*i*JO1366) and *Geobacter metallireducens* GS-15 (*i*AF987), as well as with other bacteria that also possess intracellular stages during their growth cycles, such as *Yersinia pestis* CO92 (*i*PC815), *Salmonella* enterica subsp. enterica serovar Typhimurium str. LT2 (STM_v1_0 model) and *Shigella flexneri* (*i*SF1195).

This reaction essentiality analysis showed no significant correlation between the number of essential reactions and the size of the metabolic network or the microorganism’s lifestyle. The number of essential reactions found ranged from 214 to 419, with *Y. pestis* and *P. putida* being the organisms with lower and higher number of essential reactions, respectively. Moreover, the number of these essential reactions for the δ-proteobacteria, *B. bacteriovorus* and *G. metallireducens*, account for approximately 30% of the total reactions (Table 2 and Fig S1). This rate could be related with the lack of a secondary metabolism in this bacterial group, which should be explored in depth in order to increase the computational value of the results.

**Table 2.** Reaction essentiality of the metabolic models of different bacteria

|  | *E. coli* | *P. putida* | *G. metallireducens* | *B. bacteriovorus* | *Salmonella* | *S. flexneri* | *Y. pestis* |
| --- | --- | --- | --- | --- | --- | --- | --- |
| Total reactions | 2583 | 2826 | 1285 | 1002 | 2545 | 2630 | 1961 |
| Essential reactions* | 274 | 404 | 387 | 305 | 333 | 264 | 210 |
| % of essential rxn | 10.6 | 14.6 | 30.1 | 30.2 | 13.1 | 10.3 | 10.7 |

\*Exchange reactions were excluded

The comparison of the essential reactions of the free living organism and the intracellular bacteria provided three main groups of essential reactions (exchange reactions were excluded from the comparison) (Fig 5): i) shared essential reactions between free-living and intracellular microorganisms (38 reactions), ii) free-living microorganisms’ exclusive essential reactions (27 reactions), iii) intracellular microorganisms’ exclusive essential reactions (15 reactions). Potentially, the 38 shared reactions would be part of the hypothetical essential metabolic network. Overall, the reactions found in the shared essentiality group are related with cell envelope, nucleotide, and cofactors (Table S6).

**Fig 5.**
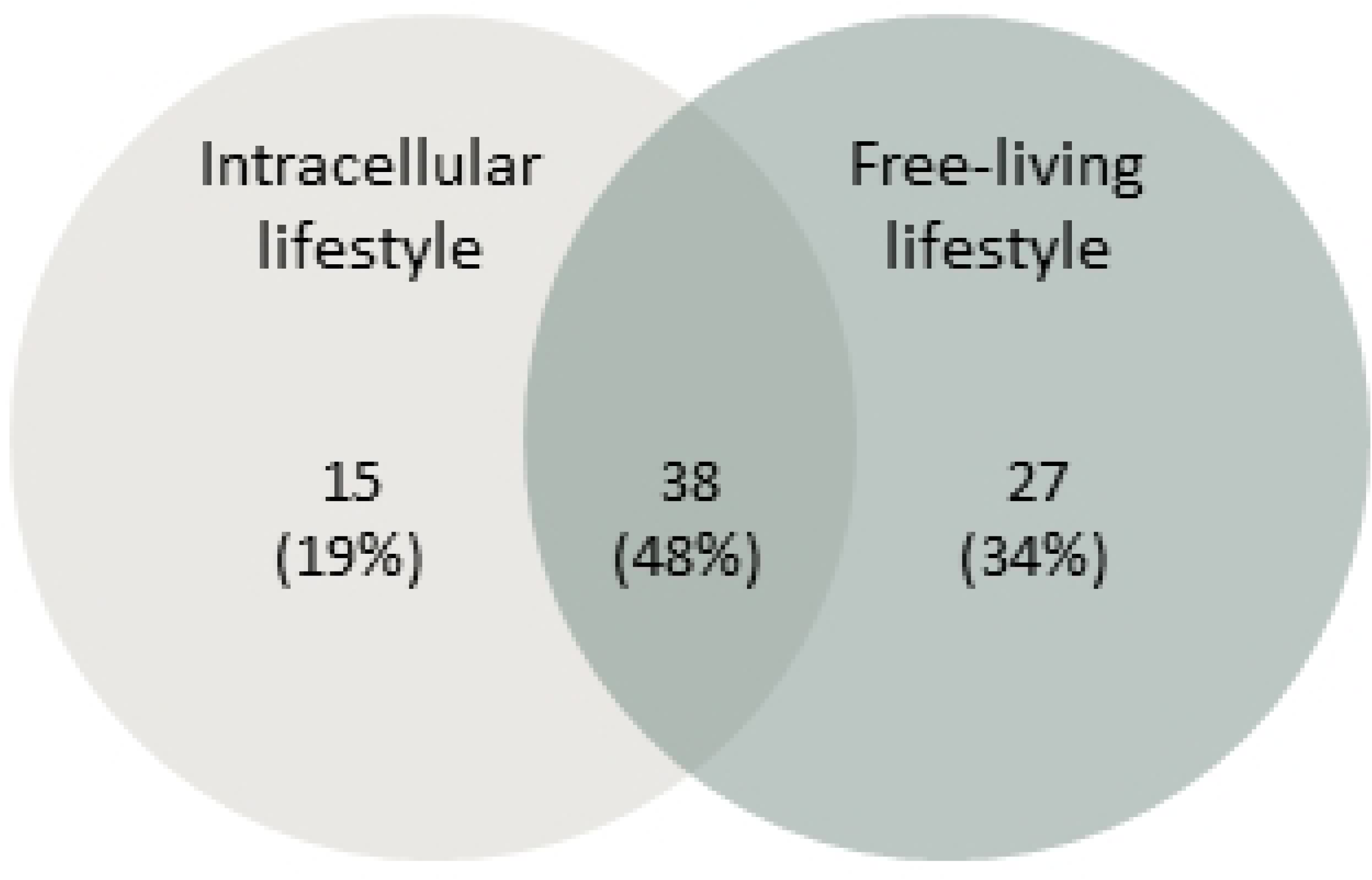
Comparison of the reaction essentiality of intracellular lifestyle and free-living bacteria.

Among the essential reactions found exclusively in the group of free-living microorganisms, none of them are present in the *i*CH457 metabolic model and they are mostly involved in cell envelope biosynthesis. This result together with the predator’s auxotrophies suggest the adaptation of *B. bacteriovorus* to a non-free-living lifestyle, where the uptake of metabolites becomes crucial to its survival. Moreover, numerous reactions involved in amino acid metabolism are included in the group of free-living organisms, but not in the predator set, likely due to the direct incorporation of these metabolites from the prey.

### Analysis of the predator’s lifestyle using condition-specific models: Attack Phase (*i*CHAP) and Growth Phase (*i*CHGP) models

*B. bacteriovorus* possesses a biphasic growth cycle, leading by an extracellular attack phase (AP) and an intraperiplasmic growth phase (GP). It has been previously reported that these two stages are clearly differentiated in terms of gene expression [49] and also in the biological role [20], changes that must be strongly determined by the microenvironment.

*B. bacteriovorus* AP cells are exposed to extracellular environment with highly diluted concentration of nutrients, but during GP the predator finds a very rich environment inside of the prey. As a consequence, it is reasonable to assume that the predator could hardly find nutrients during AP, which defines the search, attachment to and invasion of new preys as its main biological objective under this scenario. It could be envisaged that, during this period, the predator’s metabolism is rerouted to obtaining energy, in terms of ATP, thus allowing flagellum movement and facilitating the collision with prey cells thanks to its high velocity [50]. Once the predator enters the prey periplasm it uses the cytoplasm as a source of nutrients to initiate GP. Bacterial cytoplasm is a very crowded compartment where most of the components of a microorganism are localized (30-40% of macromolecules and over 70% of proteins; [51]). Thus, the cytoplasm is an extremely rich environment supporting growth and the completion of *B. bacteriovorus*’ life cycle [52, 53]. Consequently, it is reasonable to hypothesize that the main aim of this phase is to grow, which implies a highly active metabolism (catabolism and anabolism) supporting fast biomass generation. In fact, recent transcriptomic analyses have shown a highly activated anabolism in this phase [49].

To obtain a deeper understanding of this predatory bacterium in each phase of its life cycle, we constructed two different condition-specific models. Overall, these GP and AP-condition models were constructed by constraining *i*CH457 in terms of: i) nutrient availability, ii) biological objective, and iii) gene expression profile. As a first step, based on the environmental conditions, we defined two different *in silico* media: e.g., minimal and rich medium for AP and GP, respectively (Text S1). Secondly, focusing on biological role, we used different biological objectives for simulating AP and GP phases. Thus, under AP and GP, ATP production and biomass production were selected as differential objective functions, respectively. Finally, in order to constrain even more the solution space in each model, data from RNA-seq analyses collected during AP and GP [49] were integrated into the metabolic model by using GIM3E [54]. GIM3E is an algorithm which minimizes the use of reactions whose encoding gene expression levels are under a certain threshold and finds a flux distribution consistent with the target function (biomass generation for GP or ATP production for AP). Following this workflow, we constructed two new models (*i*CHAP and *i*CHGP), mimicking AP and GP growth phases, respectively (Fig 2B).

The number of reactions of each specific-condition model was significantly reduced (from 1001 to 810 and 841 in AP and GP, respectively). This significant reduction involves reduced solution spaces, and thus likely more accurate predictions. As could be inferred given the difference in biological objectives in each phase, the condition-specific models were significantly different regarding the specific metabolic content (Fig 2B and Table S7). For instance, while we found several reactions only present during AP (67 reactions), including reactions involved in glycerophospholipid degradation, the β-oxidation pathway and a large number of reactions were only present in GP model (98 reactions) i.e. reactions responsible for the biosynthesis of the cell envelope, nucleotides, fatty acids and lipids (Table S7). In other words, we found that while the unique enzymes present during AP were mainly involved in energy production and cell survival, during GP the reactions were largely involved in anabolic pathways including biosynthesis of biomass building blocks.

These reaction profiles gave way to the optimal pipeline for system exploration of the resulting solution spaces using Markov chain Monte Carlo sampling; [55] thus it was possible to establish potential differences in the metabolic states between AP and GP. Subsequently, we assessed the most probable carbon flux distribution between the two condition-specific models to reveal integrated information about the predator’s metabolism (Fig 6). Thereby the behavior during AP seems to follow a balanced oxidative metabolism aimed at energy production, including intense flux across TCA and oxidative phosphorylation. On the contrary, no significant fluxes were predicted across anaplerotic and biosynthetic pathways including gluconeogenesis, pentose phosphate, and lipid biosynthesis, which suggests negligible participation of these metabolic hubs during AP. Interestingly, a completely inverse metabolic scenario was predicted under GP. Firstly, this specific model predicted key energetic metabolic pathways being partially inactive during GP. For example, it is important to remark an incomplete performance of the TCA cycle when several stages, including citrate synthase (CS), aconitase (ACONT), isocitrate dehydrogenase (ICDH), and malate dehydrogenase (MDH) were predicted carrying no flux at all. Instead, acetyl-CoA derived from amino acid catabolism was mainly funneled to lipid biosynthesis. Reduction equivalents powering oxidative phosphorylation were produced, almost exclusively, from glutamate metabolism via α-ketoglutarate dehydrogenase and succinate dehydrogenase, thus ensuring ATP production. Finally, a very high flux through gluconeogenesis from pyruvate was predicted, thus enabling the required building blocks for nucleotide and cell envelope biosynthesis in this phase (Fig 6). Interestingly, this scenario described for GP is fully compatible with the energy-saving mechanisms suggested for *B. bacteriovours*. Therefore, the reuse of prey-derived biomass building blocks renders the role of the TCA cycle as the main source of reducing equivalents powering the production of ATP negligible.

**Fig 6.**
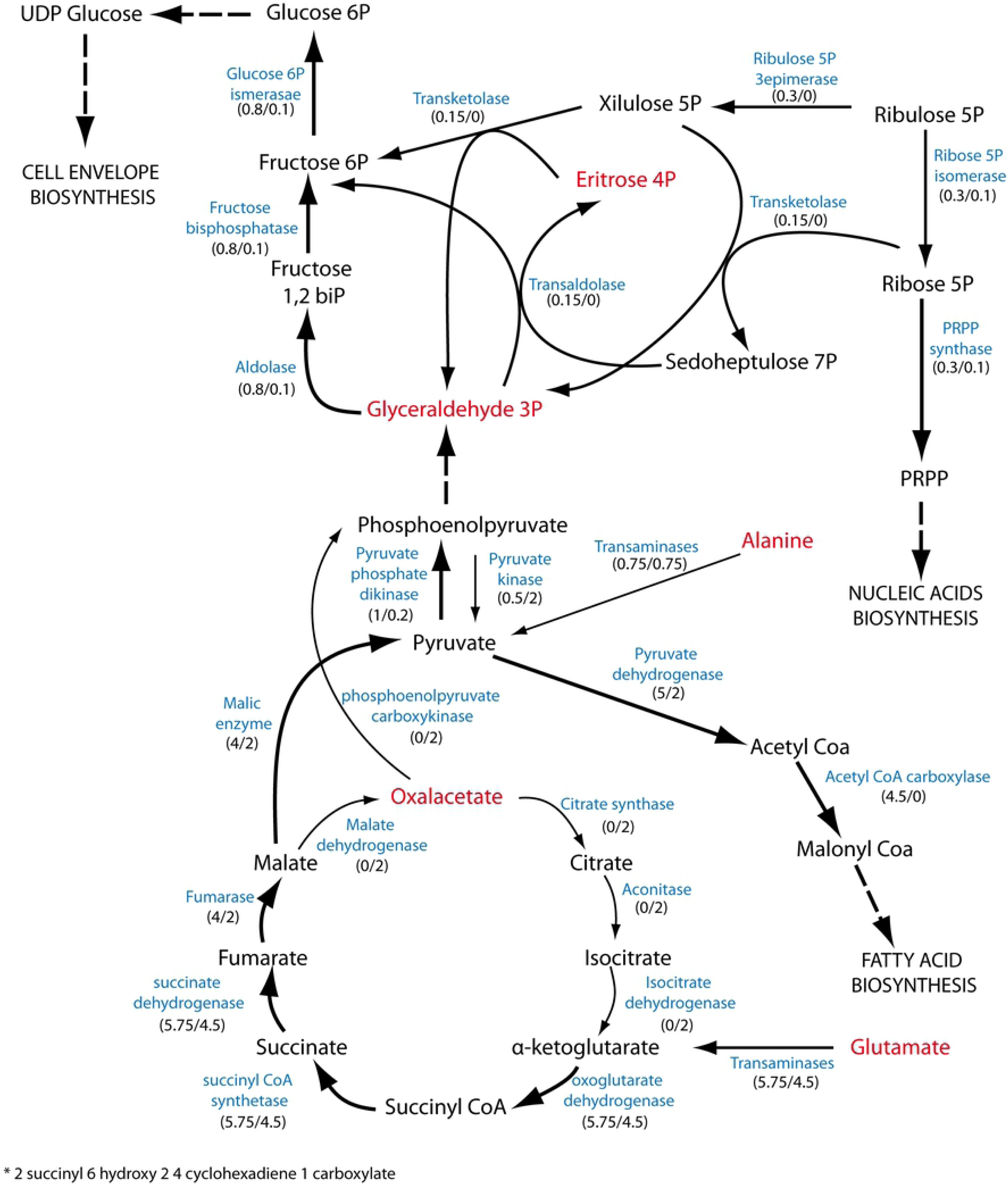
Prediction of the carbon flux distribution in *i*CH457 metabolic network. Graphical representation of the metabolic carbon fluxes during the life cycle of *B. bacteriovorus* HD100. The numbers below each reaction represent the more probable flux in each phase (GP flux/AP flux) as determined by Monte Carlo sampling analysis. The thick arrows highlight the carbon flux distribution in GP compared with AP. The thin arrows highlight the reactions that are active in AP compared with GP. As the major carbon sources are amino acids, alanine and glutamate come directly from the breakdown of the dipeptides or from the single amino acids. Eritrose 4 phosphate and glyceraldehyde 3 phosphate come from the degradation pathways of serine and threonine.

## DISCUSSION

Integrative approaches combining traditional and innovative technologies are currently being addressed to establish the metabolic network of hot-spot microorganisms. This issue becomes much more challenging when it refers to predatory microorganisms such as the bacterium *B. bacteriovorus,* which exhibit a bi-phasic lifestyle. With the aim of elucidating the metabolic network wired to predator physiology and lifestyle, we implemented a computational test-bed that proved very useful in the assessment of our predator’s phenotype-genotype relationships, while providing new insight on how *B. bacteriovorus*’ metabolism operates at the systems level.

### Complex *B. bacteriovorus* lifestyle has guided a significant genome streamlining process and the acquisition of biosynthetic energy-saving mechanisms

Comparison of the essential reactions between *B. bacteriovorus* and other intracellular lifecycle bacteria and free-living microorganisms has revealed the loss of biosynthetic pathways (Table S6, reactions exclusive to free-living microorganisms). This metabolic scenario is only possible because the host/prey metabolic machinery provides the required biomass building blocks during the intracellular stage of the growth cycle. Despite numerous auxotrophies having been reported in specific genes [6], the metabolic model has allowed the functional contextualization of these biosynthetic deficiencies within the network. For instance, model-based analyses identified additional metabolic gaps which had remained unknown so far, while on the other hand they provided alternative metabolic routes overcoming theoretical auxotrophies. Overall, our analysis has shown a significantly higher number of auxotrophies than previously thought. The loss of essential biosynthetic genes is a typical characteristic of bacteria existing in nutrient-rich environments, such as lactic acid bacteria, endosymbionts or pathogens [56]. In this sense, although *B. bacteriovorus* HD100 possesses a relatively large genome, it could also be included in this “genome streamlining” bacterial group because it directly employs whole molecules from the cytoplasm of the prey [53,57,58]. With regard to the production of the biomass building blocks, it is noteworthy that most amino acids suffer a total lack of biosynthesis pathways. In contrast, *B. bacteriovorus* is fully equipped with the biosynthetic routes for nucleotides and fatty acids. Keeping in mind the macromolecular composition of a prokaryotic cell’s cytoplasm as the natural growth niche of *B. bacteriovorus* (50 % proteins, 20 % RNA, 10 % lipids, 20 % remaining components), it is easy to speculate why the oligopeptide transporter systems are widely represented. While the factors driving *de novo* synthesis or the uptake of biomass building blocks are still unknown, it is likely that these processes are extremely regulated and only activated in the absence of intermediates. A significant flux feeding nucleic acid biosynthesis was predicted (Fig 6). Thus, an important amount of nucleotides came from *de novo* synthesis pathways. This would occur during *in vivo* conditions even in the presence of nucleotides in the extracellular medium (preýs cytoplasm). This high requirement of nucleotides beyond the amount provided by the prey could justify the presence of a complete nucleotide biosynthesis pathway in contrast with the scenario found in the biosynthesis of amino acids and cofactors when multiple autotrophies were found.

In addition, the presence of these complete metabolic pathways determines the potential ability of the predator to survive and grow without prey, as predicted by the model. Supplying the model with a rich medium based on amino acids returned a simulation which provided key information about growth and generation of biomass. Importantly, this potentially independent growth might be associated with *B. bacteriovorus*’ role as a balancer of bacterial population either in aquatic or soil environments, or in the intestine of healthy individuals, because survival of predator cells is not uniquely dependent on the predation event [48].

On the whole, our data support the hypothesis and suggest that the metabolic properties of *B. bacteriovorus* are closer to those of the postulated minimal metabolic network. This low robustness of the metabolic network suggests *Bdellovibrio* is more niche-specific than previously thought and the environmental conditions governing predation may be relatively uniform. However, in-depth studies of the metabolic capabilities of the predator are needed to complete the metabolic network and obtain more reliable *in silico* predictions.

### Nutrient availability and biological objective largely conditioned the metabolic shift from *i*CHAP to *i*CHGP

The development of *i*CH457, *i*CHAP and *i*CHGP has provided a computational framework for a better understanding of the physiological and metabolic versatility of BALOs and other predatory bacteria. In addition, it has allowed the computational demonstration, in terms of flux distribution, of the metabolic shift between the different phases. Through detailed analysis of these specific-condition models it has been possible to conclude that during AP, *B. bacteriovorus* invests most of its resources in ATP generation, presumably in order to fuel the flagellum to search for new prey. In contrast, GP was characterized by the biosynthesis of biomass building blocks. During this phase, several metabolic pathways become inactive, allowing carbon flux distribution re-routing toward biosynthetic pathways. For instance, the TCA cycle shifts from a completely operational state to an anaplerotic mode by inactivating the decarboxilative branch including citrate synthase, aconitase and isocitrate dehydrogenase. In parallel, glutamate was used as a main carbon and energy source. The metabolic switch in *B. bacteriovorus* between the different growth phases has revealed an environmental adaptation of this predator to tackle a rich medium, which would provide an explanation for the development of HI strains. Overall, the carbon flux predictions were compatible with the complex lifestyle of *Bdellovibrio* cells and provided an unprecedented overview of the metabolic shifting required to move from AP to GP, as well as new knowledge about the connections within the predator’s metabolic network.

Finally, the results obtained during this study contribute not only to increasing the available metabolic knowledge of *B. bacteriovorus*, but also to providing a computational platform for the full exploitation of this predator bacterium as a biotechnology workhorse in the near future.

## MATERIALS AND METHODS

### Genome-scale metabolic network reconstruction: *i*CH457

The genome-scale metabolic model of *B. bacterivorous* HD100 (*i*CH457) was constructed using standardized protocols for metabolic reconstruction [22, 59], and is detailed in Fig 2A. An initial draft reconstruction was generated from the annotated genome of *B. bacteriovorus* HD100 (GenBank number: BX842601.2) using the automatic application provided by Model Seed server [60]. Additionally, the metabolic content of *B. bacteriovorus* was mapped with two broadly used and high-quality GEMs belonging to *E. coli* (*i*JO1366; [61]) and *P. putida* (*i*JN1411; [46]), generating additional drafts by using MrBac Server [62]. Once these models were unified into a final reconstruction, we proceeded to a thorough manual curation of the collected metabolic information. During this iterative process, the final inclusion of each individual’s biochemical reaction was assessed using genomic [6], metabolic, transporter and GEMs databases, including: Kyoto Encyclopedia of Genes and Genomes (KEGG, [63]), BRENDA [64], BIGG [65]. Transport reactions were also added by using the TransportDB [66] database. Relevant reactions added during this process were listed in Supplementary Tables (Added reactions I and II). Finally, we performed a manual gap filling step in order to connect the network as much as possible and remove potential inconsistencies. *B. bacteriovorus* legacy literature has been thoroughly consulted, ensuring high confidence in the metabolic content included. When specific data for the HD100 strain were not available, information from phylogenetically related organisms such as 109 Davis strain was used as previously suggested [59]. The metabolites and reactions included in this metabolic model are listed in Tables S1 and S2.

### Model Analysis: Flux Balance Analysis (FBA)

FBA is by far the most popular approach for analyzing constraint-based models and it is used in many applications of GEMs. FBA uses optimization of an objective function to find a subset of optimal states in the large solution space of possible states that is shaped by the mass balance and capacity constraints. In FBA, the solution space is constrained by the statement of a steady-state, under which each internal metabolite is consumed at the same rate as it is produced [45].

The conversion into a mathematical format can be done automatically by parsing the stoichiometric coefficients from the network reaction list e.g. using the COBRA toolbox [55]. The dimensions of the stoichiometric matrix, **S**, are **m** by **n,** where **m** is the number of metabolites in the reaction network and **n** is the number of reactions. Therefore, each column represents a reaction and each row represents the stoichiometric participation of a specific metabolite in each of the reactions. FBA was used to predict growth and flux distributions. FBA is based on solving a linear optimization problem by maximizing or minimizing a given objective function to a set of constraints. The foundations and applications of FBA have been reviewed elsewhere [67, 68]. A particular flux distribution of the network, **v**, indicates the flux levels through each of the reactions. Based on the principle of conservation of mass and the assumption of a steady state, the flux distribution through a reaction network can be characterized by the following equation: **S** x **v** = 0 [45, 69]. Constraints are placed on individual reactions to establish the upper and lower bounds on the range of flux values that each of the reactions can have. These constraints are described as follows: **α_1_ ≤v_1_≤β_1_**, where **α1** is the lower bound on flux **v1**, and **β1** is the upper bound. If no information about flux levels is available, the value of **α_1_** is set to zero for irreversible fluxes. In all other cases, **α_1_** and **β_1_** are left unconstrained, thus allowing the flux to take on any value, whether positive or negative.

### Biomass function

It is commonly assumed that the objective of living organisms is to divide and proliferate. Thus, many metabolic network reconstructions have a so-called biomass function, in which all known metabolic precursors of cellular biomass are grouped (e.g. amino acids, nucleotides, phospholipids, vitamins, cofactors, energetic requirements, etc.). Since no detailed studies about *B. bacteriovorus* biomass composition are available, the biomass composition from *P. putida* [46] was used as a template for the biomass function of *i*CH457. However, data from *B. bacteriovorus* were added when available (e.g. nucleotide composition -from genome sequence). The detailed calculation of biomass composition is provided in Table S3.

### Generation of growth phase-specific models: *i*CHAP and *i*CHGP

A given metabolic reconstruction is defined by the metabolic content contained in the genome and thus is unique for the target organism. However, it is possible to construct different condition-specific models by applying additional constraints such as condition-specific data (including physiological), gen/protein expression and flux data, etc.

To construct condition-specific metabolic models we incorporated these additional constraints to the model by means of a stepwise procedure including condition specific: i) biomass, ii) nutrient availability and iii) gene expression data (Fig 2B). Firstly, the objective function was adjusted to the biological role of AP and GP. ATP maintenance and biomass equations were selected as objective functions for AP and GP, respectively. In addition, different *in silico* media were designed for each phase, simulating the availability of nutrients in each growth phase (S1 Text). Finally, available AP and GP gene expression datasets [49] were incorporated in order to constrain even further the solution space us GIM3E [70]. GIM3E builds reduced models by removing the reactions not available in the expression dataset while preserving model functionality. It should be noted that GIM3E considers which genes are expressed or not, but not the modifications in mRNA levels under different experimental conditions. A given gene was considered expressed when its RNA levels in the RNA-seq analysis fell within the first quartile, which is ≥ 10 RPKM using the available dataset [49].

The distribution of possible fluxes in the specific-condition models was calculated using Markov chain Monte Carlo sampling [55]. The median value from the distribution was used as the reference flux value.

### Reactions essentiality analysis

In order to determine the effect of a single reaction deletion, all the reactions associated with each gene in *i*CH457 were individually suppressed from the matrix ***S***. FBA was used to predict the mutation growth phenotype. The *singleReactionDeletion* function implemented in the COBRA Toolbox [55] was used to simulate knockouts. A lethal deletion was defined as that yielding < 10% of the original model’s growth rate values. The simulations for reaction essentiality were performed using the rich *in silico* medium for *i*CH457 (Supporting information). Reaction essentiality analysis has been performed for other bacteria: *P. putida* KT2440 (*i*JN1411) [46], *E. coli* strain K-12 substrain MG1655 (*i*JO1366) [71], *Geobacter metallireducens* GS-15 (*i*AF987) [72], *Yersinia pestis* CO92 (*i*PC815) [73], *Salmonella* enterica subsp. enterica serovar Typhimurium str. LT2 (STM_v1_0 model) [65] and *Shigella flexneri* (*i*SF1195) [65].

The associations between essential reactions and each bacterium were represented building a bipartite network. For visualization we use Gephi software (0.9.2). These essential reactions and the bacterial models were clustered in a heatmap using the pheatmap (v. 1.0.12) package within the R environment (http://www.R-project.org).

### Software

The *i*CH457 model was analyzed with the COBRA Toolbox v2.0 within the MATLAB environment (The MathWorks Inc.) [74]. Tomlab CPLEX and the GNU Linear Programming Kit (http://www.gnu.org/software/glpk) were used for solving the linear programing problems.

## AUTHOR CONTRIBUTION

M.A.P. and J.N. designed research. C.H. and J.N. performed the reconstruction. C.H. performed the analysis. C.H, M.A.P and J.N analyzed data. C.H and J.N drafted the paper. All the authors contributed to the final version.

## ACKNOWLEDGEMENTS

This work was supported in part by the Spanish Ministry of Economy and Competitiveness through funding provided to project BIO2014-59528-JIN, BIO2013-44878-R, Engicoin 760994 and BIO2017-83448-R.

The authors thank Clive A. Dove for critical reading of the manuscript.

## REFERENCES

1. Thompson JN. The evolution of species interactions. Science. 1999;284: 2116–8. doi:10.1126/science.284.5423.2116

2. Clarholm M. Microbes as predators or prey. Current perspectives on microbial ecology. 1984.

3. Stolp H, Starr MP. Bdellovibrio bacteriovorus gen. etsp. n., a predatory, ectoparasitic, and bacteriolytic microorganism. Antonie Van Leeuwenhoek. 1963;29: 217–248.

4. Jurkevitch E, Davidov Y. Phylogenetic Diversity and Evolution of Predatory Prokaryotes. ACS Division of Fuel Chemistry, Preprints. 2006. doi:10.1007/7171

5. Caballero JDD, Vida R, Cobo M, Máiz L, Suárez L, Galeano J, et al. Individual Patterns of Complexity in Including Predator Bacteria, over a 1-Year Period. 2017;8: e00959–17. doi:10.1128/mBio.00959-17

6. Rendulic S, Jagtap P, Rosinus A, Eppinger M, Baar C, Lanz C, et al. A predator unmasked: life cycle of Bdellovibrio bacteriovorus from a genomic perspective. Science. 2004;303: 689–92. doi:10.1126/science.1093027

7. Hobley L, Lerner TR, Williams LE, Lambert C, Till R, Milner DS, et al. Genome analysis of a simultaneously predatory and prey-independent, novel Bdellovibrio bacteriovorus from the River Tiber, supports in silico predictions of both ancient and recent lateral gene transfer from diverse bacteria. BMC Genomics. 2012;13: 670. doi:10.1186/1471-2164-13-670

8. Wurtzel O, Dori-Bachash M, Pietrokovski S, Jurkevitch E, Sorek R, Ben-Jacob E. Mutation Detection with Next-Generation Resequencing through a Mediator Genome. PLoS One. 2010;5. doi:10.1371/journal.pone.0015628

9. Sockett RE. Predatory lifestyle of Bdellovibrio bacteriovorus. Annu Rev Microbiol. 2009;63: 523–39. doi:10.1146/annurev.micro.091208.073346

10. Marta, EKSZTEJN, and, Mazal, VARON. Elongation and Cell Division in Bdellovibrio bacteriovorus. arch micorbiol. 1977;144: 175–181. doi:https://doi.org/10.1007/BF00410781

11. Capeness MJ, Lambert C, Lovering AL, Till R, Uchida K, Chaudhuri R, et al. Activity of Bdellovibrio Hit Locus Proteins, Bd0108 and Bd0109, Links Type IVa Pilus Extrusion/Retraction Status to Prey-Independent Growth Signalling. 2013; doi:10.1371/journal.pone.0079759

12. Scherff RH. Control of bacterial blight of soybean by Bdellovibrio bacteriovorus. Phytopathology. 1973;63: 400–402.

13. Loozen G, Boon N, Pauwels M, Slomka V, Rodrigues Herrero E, Quirynen M, et al. Effect of Bdellovibrio bacteriovorus HD100 on multispecies oral communities. Anaerobe. Elsevier Ltd; 2015;35: 45–53. doi:10.1016/j.anaerobe.2014.09.011

14. Atterbury RJ, Hobley L, Till R, Lambert C, Capeness MJ, Lerner TR, et al. Effects of orally administered Bdellovibrio bacteriovorus on the well-being and Salmonella colonization of young chicks. Appl Environ Microbiol. 2011;77: 5794–5803. doi:10.1128/AEM.00426-11

15. Cao H, He S, Wang H, Hou S, Lu L, Yang X. Bdellovibrios, potential biocontrol bacteria against pathogenic Aeromonas hydrophila. Vet Microbiol. Elsevier B.V.; 2012;154: 413–418. doi:10.1016/j.vetmic.2011.07.032

16. Martínez V, de la Peña F, García-Hidalgo J, de la Mata I, García JL, Prieto MA. Identification and biochemical evidence of a medium-chain-length polyhydroxyalkanoate depolymerase in the Bdellovibrio bacteriovorus predatory hydrolytic arsenal. Appl Environ Microbiol. 2012;78: 6017–26. doi:10.1128/AEM.01099-12

17. Martínez V, Herencias C, Jurkevitch E, Auxiliadora Prieto M. Engineering a predatory bacterium as a proficient killer agent for intracellular bio-products recovery: The case of the polyhydroxyalkanoates. Nat Publ Gr. 2016; doi:10.1038/srep24381

18. Margulis L. Archaeal-eubacterial mergers in the origin of Eukarya: phylogenetic classification of life. Proc Natl Acad Sci U S A. 1996;93: 1071–1076. doi:10.1073/pnas.93.3.1071

19. Jurkevitch E. Predatory Behaviors in Bacteria-Diversity and Transitions. Microbe. 2007;2: 67–73. doi:10.1128/microbe.2.67.1

20. Lambert C, Hobley L, Chang C-Y, Fenton A, Capeness M, Sockett L. A predatory patchwork: membrane and surface structures of Bdellovibrio bacteriovorus. [Internet]. Advances in microbial physiology. 2009. doi:10.1016/S0065-2911(08)00005-2

21. Bordbar A, Monk JM, King ZA, Palsson BO. Constraint-based models predict metabolic and associated cellular functions. Nat Rev Genet. 2014;15: 107–120. doi:10.1038/nrg3643

22. Monk J, Nogales J, Palsson BO. Optimizing genome-scale network reconstructions. Nature Biotechnology. 2014. doi:10.1038/nbt.2870

23. O ’brien EJ, Monk JM, Palsson BO. Using Genome-Scale Models to Predict Biological Capabilities. Cell. 2015;161: 971–987. doi:10.1016/j.cell.2015.05.019

24. Nielsen J. Systems Biology of Metabolism. Annu Rev Biochem. 2017;86: 245–275. doi:10.1146/annurev-biochem-061516-044757

25. Heirendt L, Arreckx S, Pfau T, Mendoza SN, Richelle A, Heinken A, et al. Creation and analysis of biochemical constraint-based models: the COBRA Toolbox v3.0. Nat Protoc. 2018;2: 1290–1307. doi:10.1038/nprot.2007.99

26. Lewis NE, Nagarajan H, Palsson BO. Constraining the metabolic genotype-phenotype relationship using a phylogeny of in silico methods. Nat Rev Microbiol. Nature Publishing Group; 2012;10: 291–305. doi:10.1038/nrmicro2737

27. Nguyen N-AT, Sallans L, Kaneshiro ES. The major glycerophospholipids of the predatory and parasitic bacterium Bdellovibrio bacteriovorus HID5. Lipids. 2008;43: 1053–63. doi:10.1007/s11745-008-3235-9

28. Müller FD, Beck S, Strauch E, Linscheid MW. Bacterial predators possess unique membrane lipid structures. Lipids. 2011;46: 1129–40. doi:10.1007/s11745-011-3614-5

29. Lerner TR, Lovering AL, Bui NK, Uchida K, Aizawa SI, Vollmer W, et al. Specialized peptidoglycan hydrolases sculpt the intra-bacterial niche of predatory Bdellovibrio and increase population fitness. PLoS Pathog. 2012;8. doi:10.1371/journal.ppat.1002524

30. Oberhardt MA, Palsson BØ, Papin JA. Applications of genome-scale metabolic reconstructions. Mol Syst Biol. 2009;5. doi:10.1038/msb.2009.77

31. Sun J, Sayyar B, Butler JE, Pharkya P, Fahland TR, Famili I, et al. Genome-scale constraint-based modeling of Geobacter metallireducens. BMC Syst Biol. BioMed Central; 2009;3: 15. doi:10.1186/1752-0509-3-15

32. Stolyar S, Van Dien S, Hillesland KL, Pinel N, Lie TJ, Leigh JA, et al. Metabolic modeling of a mutualistic microbial community. Mol Syst Biol. 2007;3: 92. doi:10.1038/msb4100131

33. Flowers JJ, Richards MA, Baliga N, Meyer B, Stahl DA. Constraint-based modelling captures the metabolic versatility of Desulfovibrio vulgaris. Environ Microbiol Rep. 2018;10: 190–201. doi:10.1111/1758-2229.12619

34. Hespell RB, Miozzari GF, Rittenberg SC. Ribonucleic acid destruction and synthesis during intraperiplasmic growth of Bdellovibrio bacteriovorus. J Bacteriol. 1975;123: 481–91. Available: http://www.pubmedcentral.nih.gov/articlerender.fcgi?artid=235752&tool=pmcentrez&rendertype=abstract

35. Ruby EG, McCabe JB, Barke JI. Uptake of intact nucleoside monophosphates by Bdellovibrio bacteriovorus 109J. J Bacteriol. 1985;163: 1087–94. Available: http://www.pubmedcentral.nih.gov/articlerender.fcgi?artid=219242&tool=pmcentrez&rendertype=abstract

36. Matin a, Rittenberg SC. Kinetics of deoxyribonucleic acid destruction and synthesis during growth of Bdellovibrio bacteriovorus strain 109D on pseudomonas putida and escherichia coli. J Bacteriol. 1972;111: 664–73. Available: http://www.pubmedcentral.nih.gov/articlerender.fcgi?artid=251338&tool=pmcentrez&rendertype=abstract

37. Kuenen JG, Rittenberg SC. Incorporation of long-chain fatty acids of the substrate organism by Bdellovibrio bacteriovorus during intraperiplasmic growth. J Bacteriol. 1975;121: 1145–57. Available: http://www.pubmedcentral.nih.gov/articlerender.fcgi?artid=246047&tool=pmcentrez&rendertype=abstract

38. Barabote RD, Rendulic S, Schuster SC, Saier MH. Comprehensive analysis of transport proteins encoded within the genome of Bdellovibrio bacteriovorus. Genomics. 2007;90: 424–46. doi:10.1016/j.ygeno.2007.06.002

39. Hespell RB, Rosson R a, Thomashow MF, Rittenberg SC. Respiration of Bdellovibrio bacteriovorus strain 109J and its energy substrates for intraperiplasmic growth. J Bacteriol. 1973;113: 1280–8. Available: http://www.pubmedcentral.nih.gov/articlerender.fcgi?artid=251695&tool=pmcentrez&rendertype=abstract

40. Seidler RJ, Starr M. Isolation and Characterization of Host-Independent Bdellovibrios. J Bacteriol. 1969;100: 769–785. Available: https://www.ncbi.nlm.nih.gov/pmc/articles/PMC250157/pdf/jbacter00385-0249.pdf

41. Cotter TW, Thomashow MF. Identification of a Bdellovibrio bacteriovorus genetic locus, hit, associated with the host-independent phenotype. J Bacteriol. 1992;174: 6018–24. Available: http://www.pubmedcentral.nih.gov/articlerender.fcgi?artid=207666&tool=pmcentrez&rendertype=abstract

42. Roschanski N, Klages S, Reinhardt R, Linscheid M, Strauch E. Identification of genes essential for prey-independent growth of Bdellovibrio bacteriovorus HD100. J Bacteriol. 2011;193: 1745–1756. doi:10.1128/JB.01343-10

43. Ishiguro EE. Minimum nutritional requirements for growth of host-independent derivatives of Bdellovibrio bacteriovorus strain 109 Davis. Can J Microbiol. 1974;20: 263–265. doi:10.1139/m74-041

44. Fong SS, Joyce AR, Palsson BØ. Parallel adaptive evolution cultures of Escherichia coli lead to convergent growth phenotypes with different gene expression states. Genome Res. 2005;15: 1365–1372. Available: https://www.ncbi.nlm.nih.gov/pmc/articles/PMC1240078/pdf/00151365.pdf

45. Orth JD. What is flux balance analysis? Nat Biotechnol. 2010;28: 245–248. doi:doi:10.1038/nbt.1614

46. Nogales J, Gudmundsson S, Duque E, Luis Ramos J, Putida P. Expanding the computable reactome in Pseudomonas putida reveals metabolic cycles providing robustness. 2 3 Running Head: Metabolic Robustness Cycles in. BioRxiv. 2017;May. doi:10.1101/139121

47. Seidler RJ, Starr MP, Mandel M. Deoxyribonucleic acid characterization of Bdellovibrios. J Bacteriol. 1969;100: 786–90. Available: http://www.pubmedcentral.nih.gov/articlerender.fcgi?artid=250158&tool=pmcentrez&rendertype=abstract

48. Dwidar M, Im H, Seo JK, Mitchell RJ. Attack-Phase Bdellovibrio bacteriovorus Responses to Extracellular Nutrients Are Analogous to Those Seen During Late Intraperiplasmic Growth. Microb Ecol. 2017; doi:10.1007/s00248-017-1003-1

49. Karunker I, Rotem O, Dori-Bachash M, Jurkevitch E, Sorek R. A Global Transcriptional Switch between the Attack and Growth Forms of Bdellovibrio bacteriovorus. PLoS One. 2013;8: e61850. doi:10.1371/journal.pone.0061850

50. Erkin Kuru, 1 7, Carey Lambert, 2, Jonathan Rittichier, 3 7, et al. Fluorescent D-amino-acids reveal bi-cellular cell wall modifications important for Bdellovibrio bacteriovorus predation. Nat Microbiol. 2017; doi:doi: 10.1038/s41564-017-0029-y

51. Zimmerman SB, Trach S 0. Estimation of Macromolecule Concentrations and Excluded Volume Effects for the Cytoplasm of Escherichia coli. J Mol Bid. 1991;222: 599–620. Available: http://ac.els-cdn.com/002228369190499V/1-s2.0-002228369190499V-main.pdf?_tid=fe412f06-55c4-11e7-8e6b-00000aab0f27&acdnat=1497969202_40de091c7e85b960895c477a2b0bfa60

52. Zhang Y-HP. Substrate channeling and enzyme complexes for biotechnological applications. Biotechnol Adv. 2011;29: 715–725. doi:10.1016/j.biotechadv.2011.05.020

53. Vendeville A s, Lariviere D, Fourmentin E. An inventory of the bacterial macromolecular components and their spatial organization’. FEMS Microbiol Rev. 2011;35: 395–414. doi:10.1111/j.1574-6976.2010.00254.x

54. Becker SA, Palsson BO. Context-specific metabolic networks are consistent with experiments. PLoS Comput Biol. 2008;4. doi:10.1371/journal.pcbi.1000082

55. Schellenberger J, Que R, Fleming RMT, Thiele I, Orth JD, Feist AM, et al. Quantitative prediction of cellular metabolism with constraint-based models: The COBRA Toolbox v2.0. Nat Protoc. 2011;6: 1290–1307. doi:10.1038/nprot.2011.308

56. D’Souza G, Waschina S, Pande S, Bohl K, Kaleta C, Kost C. LESS IS MORE: SELECTIVE ADVANTAGES CAN EXPLAIN THE PREVALENT LOSS OF BIOSYNTHETIC GENES IN BACTERIA. Evolution (N Y). 2014;68: 2559–2570. doi:doi:10.1111/evo.12468

57. Rittenberg SC, Langley D. Utilization of nucleoside monophosphates per Se for intraperiplasmic growth of Bdellovibrio bacteriovorus. J Bacteriol. 1975;121: 1137–44. Available: http://www.pubmedcentral.nih.gov/articlerender.fcgi?artid=246046&tool=pmcentrez&rendertype=abstract

58. Golding I, Cox EC. Physical nature of bacterial cytoplasm. Phys Rev Lett. 2006;96: 14–17. doi:10.1103/PhysRevLett.96.098102

59. Thiele I, Palsson B. A protocol for generating a high-quality genome-scale metabolic reconstruction. Nat Protoc. 2010;5: 93–121. doi:10.1038/nprot.2009.203.A

60. Aziz RK, Devoid S, Disz T, Edwards RA, Henry CS, Olsen GJ, et al. SEED Servers: High-Performance Access to the SEED Genomes, Annotations, and Metabolic Models. PLoS One. 2012;7: 1–10. doi:10.1371/journal.pone.0048053

61. Feist AM, Henry CS, Reed JL, Krummenacker M, Joyce AR, Karp PD, et al. A genome-scale metabolic reconstruction for Escherichia coli K-12 MG1655 that accounts for 1260 ORFs and thermodynamic information. Mol Syst Biol. 2007;3: 121. doi:10.1038/msb4100155

62. Liao YC, Huang TW, Chen FC, Charusanti P, Hong JSJ, Chang HY, et al. An experimentally validated genome-scale metabolic reconstruction of Klebsiella pneumoniae MGH 78578, iYL1228. J Bacteriol. 2011;193: 1710–1717. doi:10.1128/JB.01218-10

63. Ogata H, Goto S, Sato K, Fujibuchi W, Bono H, Kanehisa M. KEGG: Kyoto encyclopedia of genes and genomes. Nucleic Acids Res. 1999;27: 29–34. doi:10.1093/nar/27.1.29

64. Placzek S, Schomburg I, Chang A, Jeske L, Ulbrich M, Tillack J, et al. BRENDA in 2017: New perspectives and new tools in BRENDA. Nucleic Acids Res. 2017;45: D380–D388. doi:10.1093/nar/gkw952

65. King ZA, Lu J, Dräger A, Miller P, Federowicz S, Lerman JA, et al. BiGG Models: A platform for integrating, standardizing and sharing genome-scale models. Nucleic Acids Res. 2016;44: D515–D522. doi:10.1093/nar/gkv1049

66. Elbourne LDH, Tetu SG, Hassan KA, Paulsen IT. TransportDB 2.0: A database for exploring membrane transporters in sequenced genomes from all domains of life. Nucleic Acids Res. 2017;45: D320–D324. doi:10.1093/nar/gkw1068

67. Bonarius HPJ, Schmid G, Tramper J. Flux analysis of underdetermined metabolic networks: The quest for the missing constraints. Trends Biotechnol. 1997;15: 308–314. doi:10.1016/S0167-7799(97)01067-6

68. Varma A, Palsson B 0. Stoichiometric Flux Balance Models Quantitatively Predict Growth and Metabolic By-Product Secretion in Wild-Type Escherichia coli W3110. Appl Environ Microbiol. 1994; 3724–3731. Available: http://cepac.cheme.cmu.edu/pasilectures/costas/varma-palsson94.pdf

69. Varma A, Palsson B. Metabolic Capabilities of Escherichia coli: I. Synthesis of biosynthetic precursors and cofactors. Journal Theor Biol. 1993;165: 477–502. Available: https://ac.els-cdn.com/S0022519383712026/1-s2.0-S0022519383712026-main.pdf?_tid=f7a7d5dc-6338-4d72-9aa4-58e94bb72d4a&acdnat=1521540528_c1a58e5d5350a4be70af328083507e3f

70. Nogales J, Agudo L. A Practical Protocol for Integration of Transcriptomics Data into Genome-Scale Metabolic Reconstructions. Hydrocarb Lipid Microbiol Protoc - Springer Protoc Handbooks. 2015; 135–152. doi:10.1007/8623

71. Orth JD, Conrad TM, Na J, Lerman JA, Nam H, Feist AM, et al. A comprehensive genome-scale reconstruction of Escherichia coli metabolism—2011. Mol Syst Biol. 2011;7: 535. doi:10.1038/msb.2011.65

72. Feist AM, Nagarajan H, Rotaru AE, Tremblay PL, Zhang T, Nevin KP, et al. Constraint-Based Modeling of Carbon Fixation and the Energetics of Electron Transfer in Geobacter metallireducens. PLoS Comput Biol. 2014;10: 1–10. doi:10.1371/journal.pcbi.1003575

73. Charusanti P, Chauhan S, McAteer K, Lerman JA, Hyduke DR, Motin VL, et al. An experimentally-supported genome-scale metabolic network reconstruction for Yersinia pestis CO92. BMC Syst Biol. BioMed Central Ltd; 2011;5: 163. doi:10.1186/1752-0509-5-163

74. Hyduke D, Schellenberger J, Que R, Fleming R, Thiele I, Orth J, et al. PROTOCOL EXCHANGE | COMMUNITY CONTRIBUTED COBRA Toolbox 2. 0. Protoc Exch. 2011; 1–30.

